# Autofluorescence-based Label-free Cell Counting Method in Suspension Culture with Microcarriers

**DOI:** 10.1101/2025.09.16.676704

**Authors:** Takashi Morikura, Katsuhisa Sakaguchi, Ryu-ichiro Tanaka, Kiyotaka Iwasaki, Tatsuya Shimizu

**Affiliations:** Faculty of Science and Engineering, Waseda Research Institute for Science and Engineering, Waseda University, Tokyo, Japan; Graduate School of Science and Technology, Keio University, Kanagawa, Japan; Department of Medical Engineering, Faculty of Science and Engineering, Tokyo City University, Tokyo, Japan; Institute of Advanced Biomedical Engineering and Sciences, TWIns, Tokyo Women’s Medical University, Tokyo, Japan; Department of Modern Mechanical Engineering, Waseda University, Tokyo, Japan

## Abstract

To advance the industrialization of cultured meat and regenerative medicine, scalable and efficient cell culture techniques are essential. Among these, the suspension culture method using microcarriers has emerged as a promising approach for the large-scale cell culture technique. However, monitoring cell growth on the microcarriers remains challenging, particularly in developing cell counting techniques that can be seamlessly integrated into bioprocess workflows without cell detachment, fluorescence labeling and any parameter tuning in the analysis algorithm. In this study, we proposed a versatile image analysis-based cell counting method by using cellular autofluorescence without any parameter tuning. The proposed method estimates the number of cells by applying spatiotemporal averaging to the autofluorescence signals in the microscopic images. Using numerical and cell culture experiments, we demonstrated that the proposed method can estimates the number of cells accurately. This technique, which harnesses the ubiquitous autofluorescence inherent in living cells, offers a cost-effective and practical solution applicable to a broad range of fields requiring high-throughput cell quantification.

## Introduction

Suspension culture method using microcarriers [1, 2] is a powerful technique for large-scale cell culture in industrial biotechnology filed such as cultured meat [3–5] and regenerative medicine [6]. For the suspension culture, it is crucial to monitor the number of cells on the microcarriers over time and to perform passaging at appropriate intervals.

As the golden standard for measuring the number of cells adhering to microcarriers, protocols involving detachment and sampling of cells from the microcarriers through trypsin or accutase treatment and then counting the number of cells using a hemocytometer are commonly used [7, 8]. Fluorescent labeling techniques, which involve counting the number of fluorescently stained cells using a fluorophotometer are also widely adopted [9, 10]. However, these techniques are not suitable for time-course measurements, as the gold standard method requires cell detachment, and fluorescent labeling introduces cytotoxicity [11, 12]. Moreover, the sampling process in the golden standard method carries the risk of contamination.

To overcome these problems, alternative cell counting methods that do not require the cell detachment or fluorescence labeling have been developed, such as electrochemical methods using frequency response analyzers [13–15] and spectroscopic methods using optical spectrum analyzers [16, 17]. Nonetheless, the use of such specialized and costly equipment poses a barrier to the seamless integration into the bioprocesses and contributes to increased overall process costs.

As an alternative approach, image analysis-based method has also been developed [18, 19]. However, these previous studies are based on traditional image processing techniques, which require manual parameter tuning by experts. Versatile image analysis-based cell counting method that do not depend on such manual parameter tuning are strongly required.

In this study, we focused on an autofluorescence signal that occurs upon excitation and is widely recognized as a natural biological label [20–23]. Although autofluorescence is typically regarded as noise that reduces the signal-to-noise ratio in fluorescence staining [24, 25], we took the opposite approach. In this study, we treated autofluorescence not as noise but as a cell-specific signal. Autofluorescence is a physical phenomenon unique to the biological structure of cells; thus, we hypothesized that information regarding the number of cells could be extracted from this signal.

However, polystyrene, a common component of microcarriers, also exhibits photoluminescence [26, 27], which can obscure the autofluorescence signal emitted by cells. To address this issue, we focused on the composition of the fluorescence signal observed in microscopic images, which consists of cellular autofluorescence and photoluminescence from microcarriers. A critical assumption here is that the signals originating from the cells and from the microcarriers are generated independently and combined linearly to form the observed fluorescence signal. Assuming that these signals follow normal distributions, the fluorescence signal itself also follows a normal distribution due to the reproductive property of the normal distribution. When the autofluorescence signal from cells is relatively stronger than the microcarrier-derived signal, the mean of the fluorescence signal asymptotically approaches the mean of the cell-derived signal. Therefore, based on the law of large numbers, it may be possible to asymptotically approximate the average fluorescence signal to the mean autofluorescence signal from the cells by performing sufficiently extensive sampling. We therefore hypothesized that a cell counting method requiring no parameter tuning could be achieved by constructing an image processing algorithm based solely on the spatiotemporal averaging of autofluorescence images.

Here, we developed a versatile cell counting method by using cellular autofluorescence-based image analysis that does not require any parameter tuning. The proposed method estimates the number of cells by applying spatiotemporal averaging to the autofluorescence signals in the microscopic images. The number of cells on the microcarriers is measured using linear regression based on the spatiotemporal average values.

## Results

### Validity assessment of the proposed cell counting method based on numerical experiments

The proposed method was designed to measure the number of cells on a microcarrier by extracting cellular autofluorescence signals from the fluorescent images using spatiotemporal averaging (Fig. 1). To validate the performance of the proposed method, we performed numerical experiments based on Monte Carlo simulations

**Figure 1.**
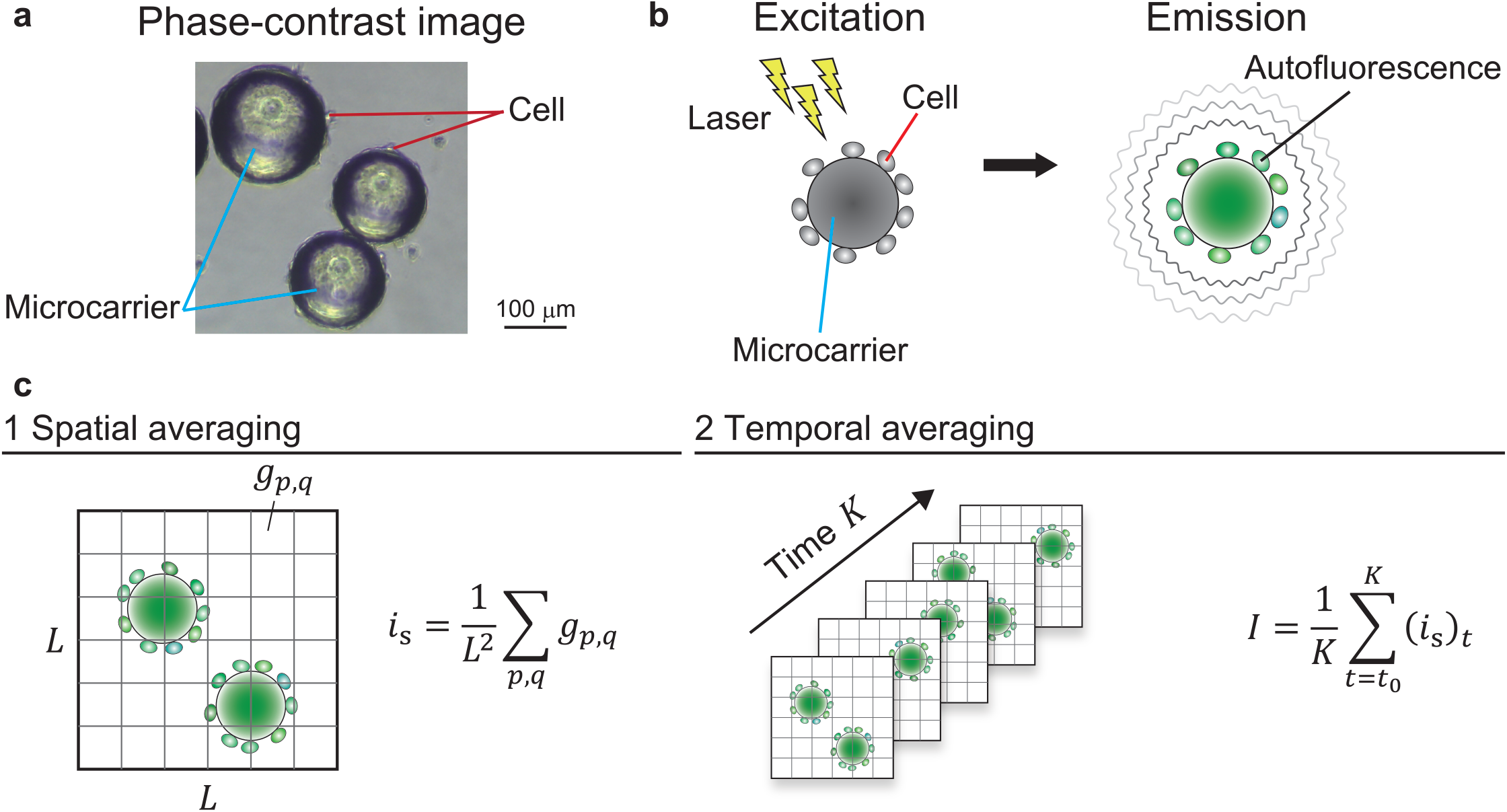
Schematic diagram of our proposed cell counting method. (**a**) Representative phase-contrast micrograph. Cells adhere to and proliferate on the surface of microcarriers. (**b**) Diagram of the autofluorescence of cells and microcarriers. When cells on the microcarriers are irradiated with excitation light, they emit autofluorescence. (**c**) Conceptual diagram of our proposed method. The spatial averaging values in gray on the fluorescence micrograph are first calculated. After acquisition of time-series data of the images, spatiotemporal average values are calculated through temporal averaging using the spatial averaging values. L indicates length of one side of the test region, *g*_*p,q*_ indicates the pixel-value of the fluorescence images, *p* and *q* indicate the spatial index in the test region. As with the equation (1) of the Supplementary Note 1, *i*_*s*_ indicates spatial average of pixel values at each time. *K* and *t*_0_ indicate the number of shots and initial time, respectively.

Figure 2 shows representative scatter plots of cell counting against the fluorescence intensity ratio and Figure 3 shows the performance of the cell counting against the fluorescence intensity ratio. As shown in Figure 2, linear coupling of the cell count to the spatiotemporal averaging value increases when the fluorescence intensity ratio increases. The result qualitatively indicates that the number of cells can be measured linearly using the spatiotemporal averaging value. The correlation coefficient and coefficient of determination increased monotonically with the fluorescence intensity ratio (Figs. 3a and 3b), and the mean absolute error and root mean square error decreased monotonically (Figs. 3c and 3d). These results suggested that the proposed method is valid if the fluorescence intensity ratio is high exists.

**Figure 2.**
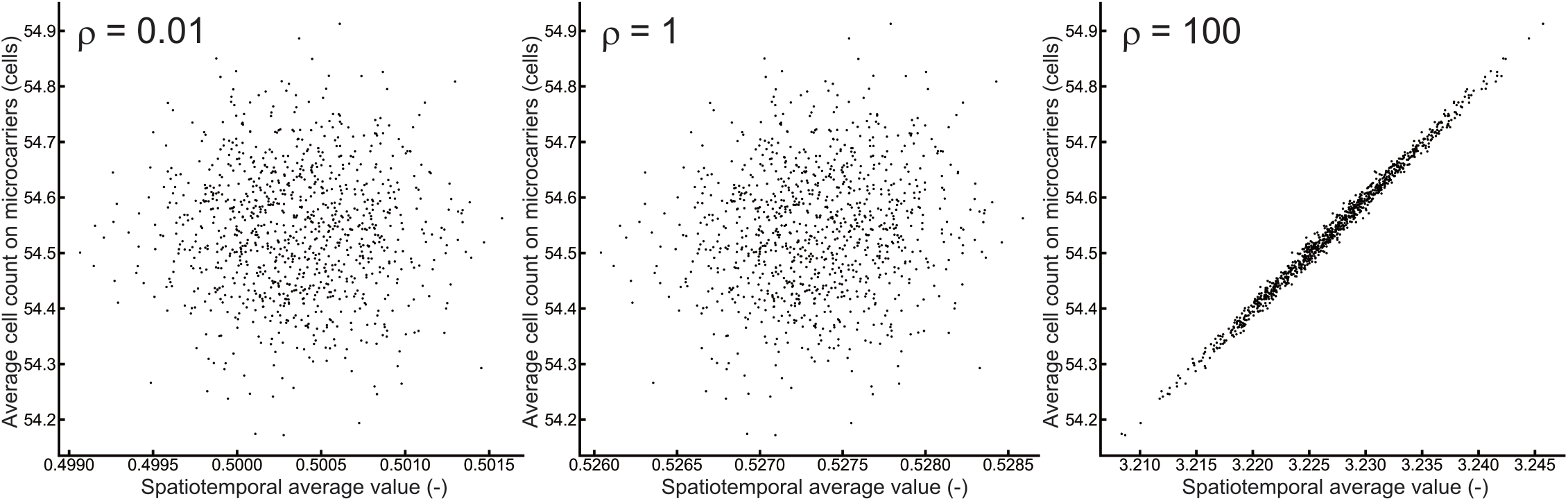
Representative scatter plots of the average cell count on microcarriers and the spatiotemporal average value. The ratio of fluorescence intensity is indicated by ρ. Dots in the figures indicate samples generated using the Monte Carlo method. The number of samples was 1000.

**Figure 3.**
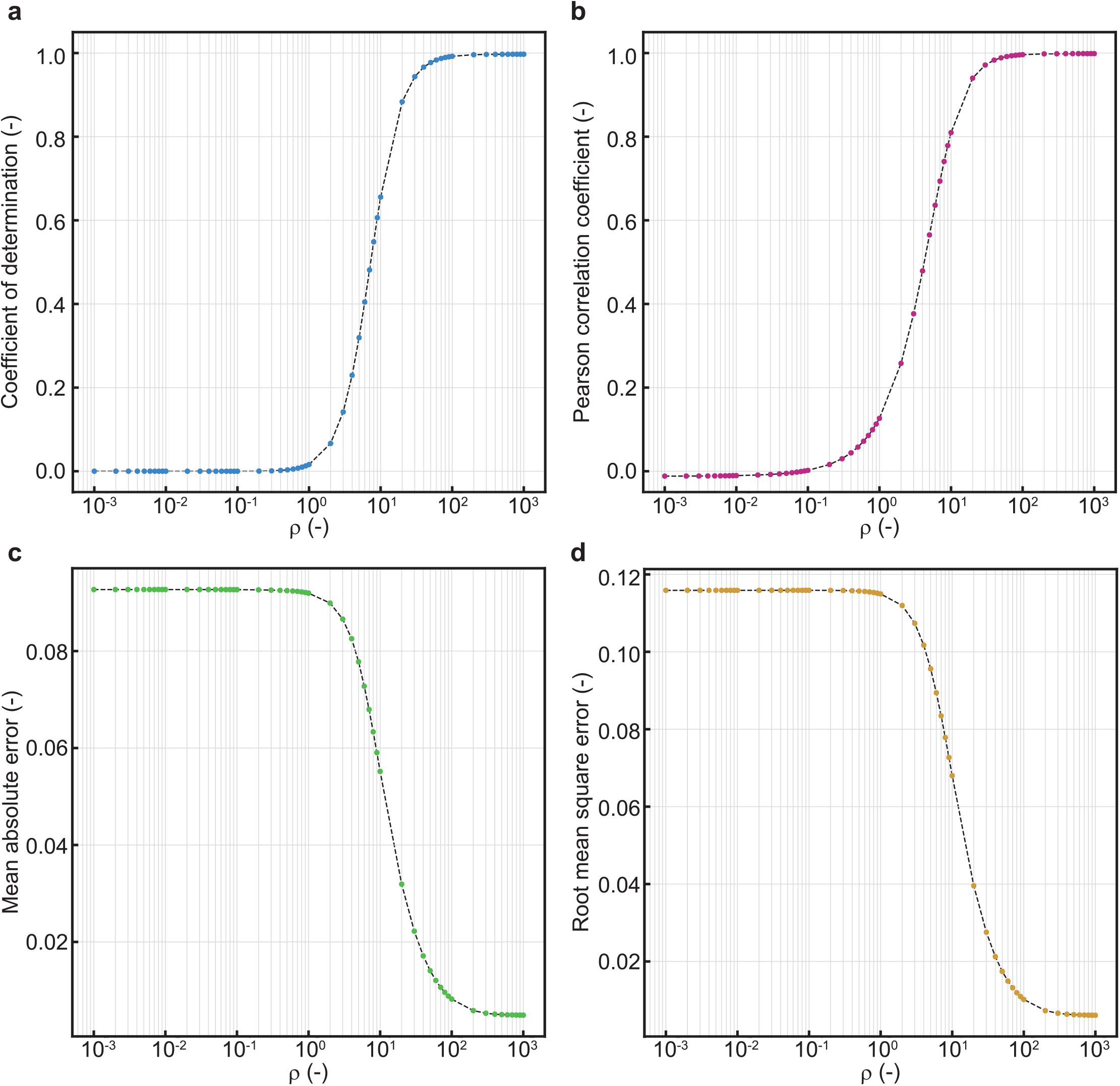
Results of numerical experiments on performance of the proposed method for the ratio of fluorescence intensity *ρ*. (**a**) Coefficient of determination. (**b**) Pearson correlation coefficient. (**c**) Mean absolute error. (**d**) Root mean square error. Dots in the figures indicate samples generated using the Monte Carlo method. The number of samples was 1000.

### Validity assessment of the proposed cell counting method based on cell culture experiments

To validate the proposed method experimentally, cell culture experiments using primary bovine myoblasts cells adhered to polystyrene spherical microcarriers were conducted. The correct number of cells as a ground truth was measured by a golden standard method using trypsin treatment and centrifugation. For the excitation of cellular autofluorescence, DAPI (excitation: 371–409 nm, emission: 430–520 nm), FITC (excitation: 450–490 nm, emission: 507–562 nm), and TRITC (excitation: 525–575 nm, emission: 593–668 nm) filters were used.

Representative fluorescence micrographs for each filter are shown in Figure 4a, and representative scatter plots are shown in Figure 4b. In the DAPI filter, only microcarriers showed strong fluorescence, whereas in the FITC filter, autofluorescence of cells on microcarriers was observed (Fig. 4a). The TRITC filter revealed weak fluorescence for both microcarriers and cells. In the scatter plots, there was no significant correlation between the spatiotemporal average value and the number of cells for DAPI and TRITC filters, whereas there was a significant correlation for the FITC filter (Fig. 4b). The coefficient of determination using the FITC filter was 0.724, mean absolute error 1.66 × 10^4^ cells/mL, and root mean square error 2.40 × 10^4^ cells/mL. Considering the count resolution of the golden standard method is in the order of 10^4^ cells/mL, these results indicated that the proposed method using the FITC filter is useful for cell counting as well as the golden standard method.

**Figure 4.**
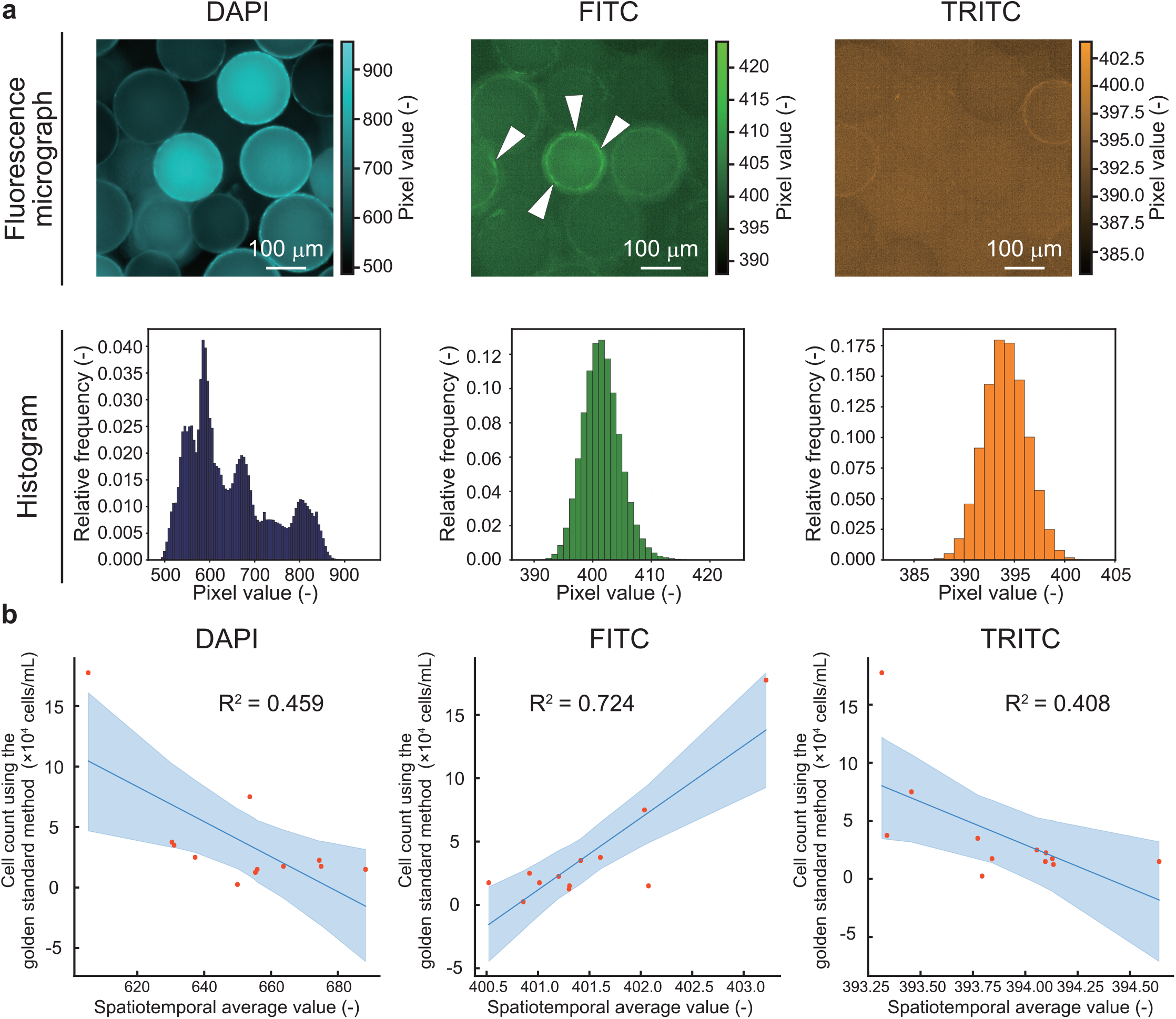
Representative results of the cell culture experiments measured by our proposed cell counting method. (**a**) Representative fluorescence micrographs and histograms of the pixel value captured by the DAPI, FITC, and TRITC filters. Relative frequency was calculated by normalizing the sum of the frequencies in each filter to 1. Although raw images were grayscale, they were colored using custom Python scripts. Blue objects in the image captured with the DAPI filter indicate the microcarriers that fluoresce due to the photoluminescence. Green objects in the images captured with the FITC filter indicate the microcarriers and cells that fluoresce due to the autofluorescence. White arrows in the image captured by the FITC filter indicates the indicate characteristic signals seem to be based on the cell-derived autofluorescence. Images captured with the TRITC filter were dominated by noise. (**b**) Representative scatter plots of our proposed method and the golden standard method. Randomly sampled sixty images were used to analyze by the proposed method. Bayesian linear regression was performed using No-U-turn sampler. Orange dots indicate values measured using our proposed method with each filter. Blue lines indicate the regression line with 95% confidential interval. The R-hat values at the end of the Markov chain Monte Carlo simulation were all 1.00, indicating convergence sufficiently. Coefficients of determination were 0.459, 0.724, and 0.408 for the DAPI, FITC, and TRITC filters, respectively. Pearson correlation coefficients were −0.677, 0.851, and −0.640 for the DAPI, FITC, and TRITC filters, respectively. The results of the statistical test for no correlation, with the significance level set at 0.01, had p-values of 0.0155, 0.000450, and 0.0250 for DAPI, FITC, and TRITC filters, respectively.

### Performance convergence and the role of the law of large numbers in the proposed method

Through the previous numerical and cell culture experiments, we demonstrated that the proposed method enables accurate estimation of cell numbers. In this section, we evaluated performance convergence and the role of the law of large numbers in the proposed method. If the proposed method operates in accordance with the law of large numbers, its performance is expected to improve asymptotically with an increasing number of samples, particularly the number of shots. To evaluate the effect of the number of shots on the performance of the proposed method, we conducted the numerical experiments and cell culture experiments.

As a result of numerical experiments, figure 5 shows performance changes of the proposed method according to the number of shots. The mean absolute error (Fig. 5a) and root-mean-square error (Fig. 5b) decreased monotonically with increasing number of images for all fluorescence intensity ratios. These results suggested that the spatiotemporal averaging value was more correlated to the number of cells with the number of shots increase.

**Figure 5.**
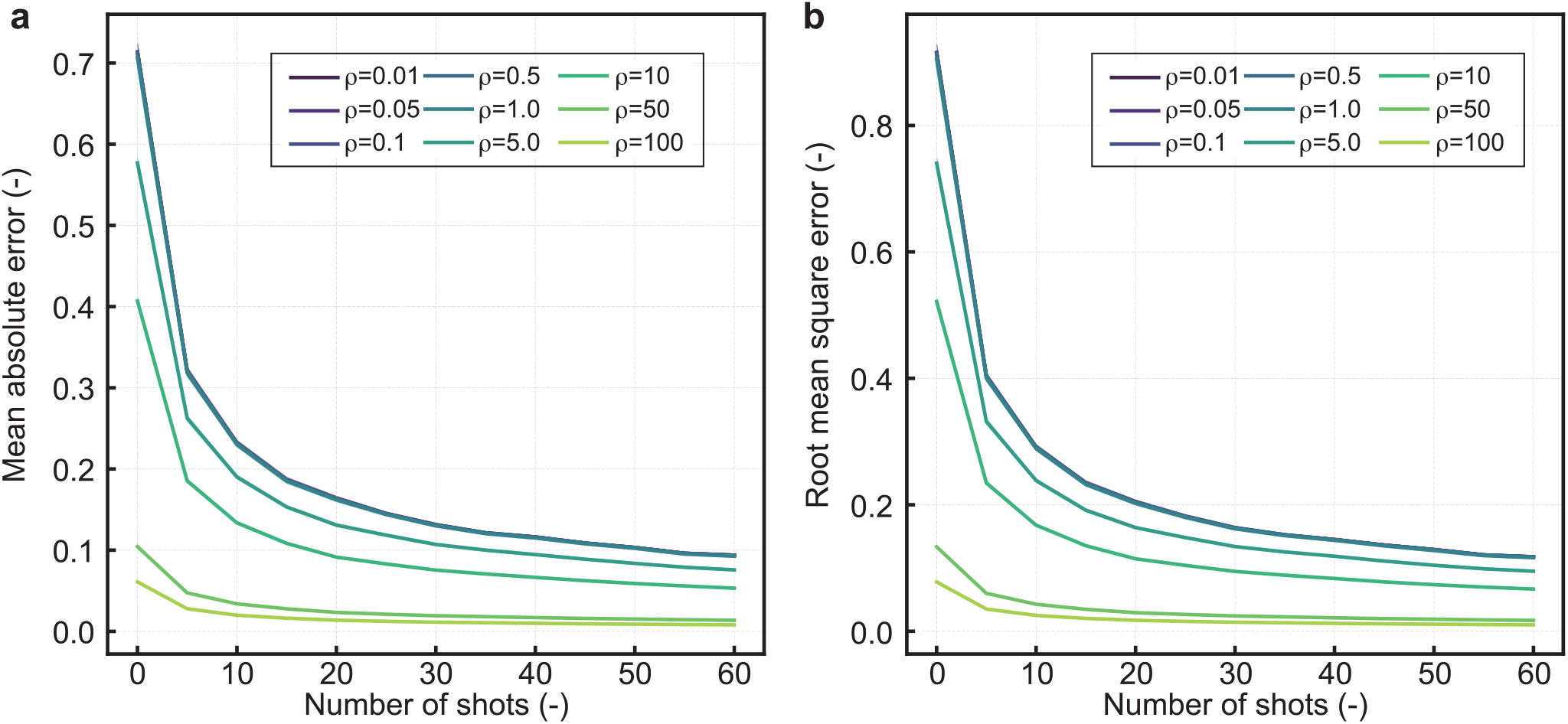
Results of numerical experiments on performance of the proposed cell counting method for the number of shots. (**a**) Mean absolute error. (**b**) Root mean square error. The ratio of fluorescence intensity is indicated by ρ. Samples were generated using the Monte Carlo method. The sample size and number of samples were 100 (mean ± 95% CI).

As a result of cell culture experiments, figure 6 shows the effects of the number of shots on the performance of the proposed method for each filter. The results of the root mean square error and absolute mean error in the cell culture experiment were almost consistent with those of the numerical experiment (Figs. 5 and 6). The reduction in these errors may be a result of following the law of large numbers, as hypothesized. The mean absolute error decreased monotonically with the number of shots for the FITC and TRITC filters, whereas it achieved steady state for the DAPI filter (Fig. 6a). The mean absolute error for the FITC filter decreased with respect to the other filters. The root mean square error decreased monotonically with the number of shots for all filters, and the root mean square error for the FITC filter decreased compared to those of the other filters (Fig. 6b). These results indicated that the performance using the FITC filter improved by increasing the number of shots better than the other filters. This finding suggested that the spatiotemporal averaging value calculated using the FITC filter was more correlated to the number of cells with the number of shots increase.

**Figure 6.**
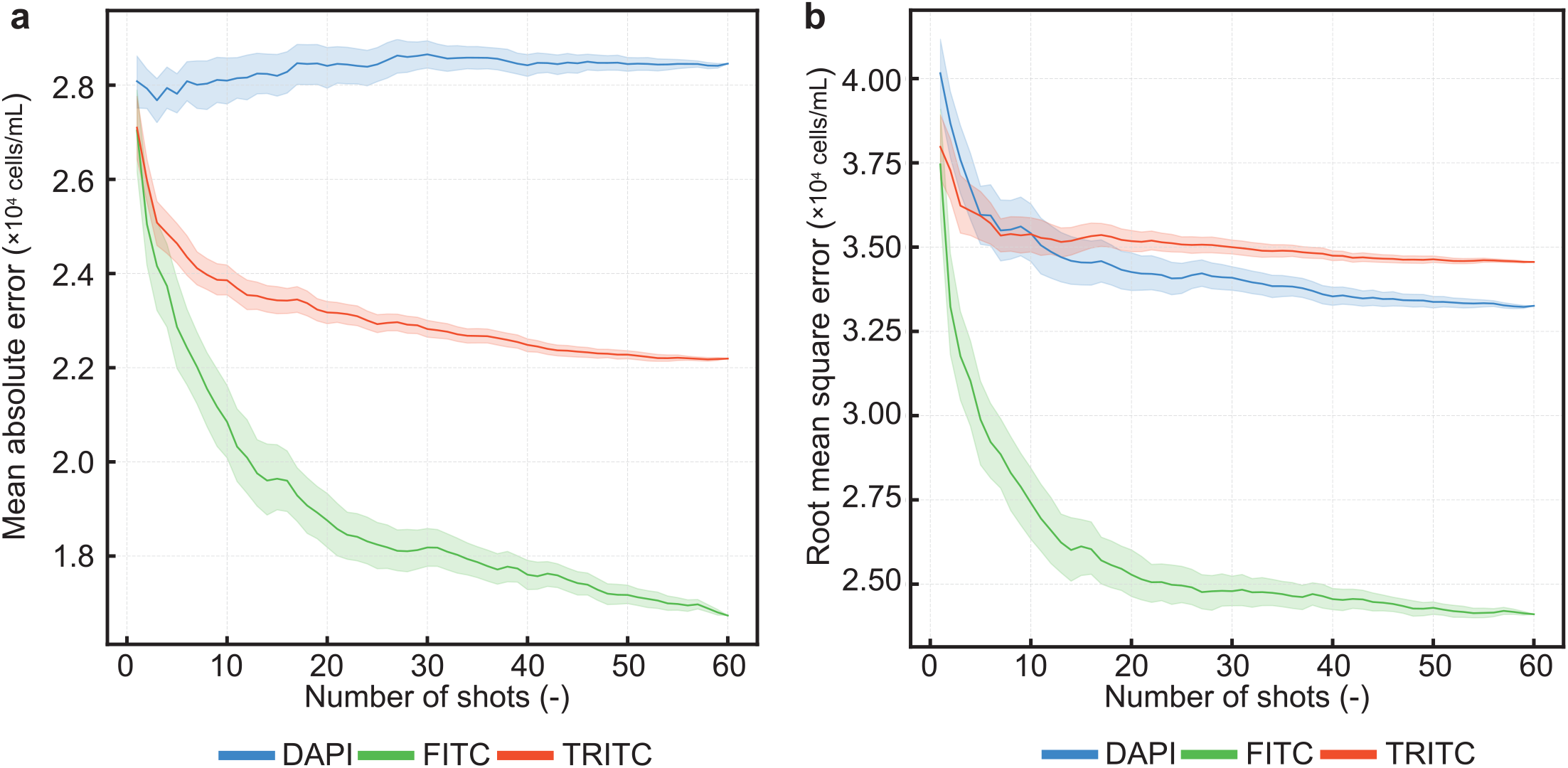
Evaluation of performance of the proposed cell counting method using DAPI, FITC, and TRITC filters. (**a**) Mean absolute error. (**b**) Root mean square error. The number of iterations for the bootstrap sampling was set to sixty (mean ± 95% CI). Blue, green, and red lines indicate the values measured with the DAPI, FITC, and TRITC filters, respectively.

## Discussion

To realize regenerative medicine, cultured meat and protein biosynthesis from cells technology at the industrial level, mass culture technology is indispensable. For mass cell culture of the anchorage-dependent proliferating cells, the suspension culture system using microcarriers has gained increasing attention. To monitoring the number of cells on the microcarriers, low-cost label-free cell counting techniques without expensive equipment are strongly desired. In this study, we developed an image analysis-based low-cost label-free cell counting method using cellular autofluorescence (Fig. 1).

Considering the results of numerical simulations (Figs. 2 and 3) and cell culture experiments (Fig. 4), it was suggested that selecting an appropriate filter based on the fluorescence wavelength ratio between cells and microcarriers is crucial for the proposed method to function effectively. Polystyrene, a component of Synthemax II, was reported to exhibit strong fluorescence from 300−400 nm [28]. In fact, the average pixel values, contrast, and contrast ratios of the Synthemax II fluorescence images were higher in the order of DAPI, FITC, and TRITC filters (see Supplementary Fig. S1). On the contrary, it was reported that the autofluorescence of cells was in the wavelength range of 450−525 nm [29]. Considering the difference in these wavelength ranges, the reason for success of the measurement using our proposed method with an FITC filter might be because of the increase in fluorescence intensity ratio of cells to microcarriers in the FITC wavelength range. Meanwhile, the low fluorescence intensity ratio for the DAPI filter and low absolute fluorescence intensity for the TRITC filter made it difficult to measure the number of cells using the proposed method.

Versatility of the proposed method is a notable characteristic. First, unlike conventional image processing algorithms that require manual parameter tunings by skilled experts or deep learning models that involve extensive hyperparameter optimization, the image analysis algorithm in the proposed method is entirely parameter-free, relying solely on simple spatiotemporal averaging. Except for the appropriate selection of a fluorescence filter, it is broadly applicable to various cell types and microcarrier materials. In addition, Second, the proposed method utilizes autofluorescence signals, which are expressed in a broad range of cell types. The most common substances that exhibit autofluorescence are NADH and FAD, which are involved in the production of ATP in the electron transport chain [29]. These substances are universally present in cells; thus, in principle, almost every cell type can be counted using our proposed method. However, the most effective wavelength region where fluorescence intensity ratio is highest may not be included in the FITC filter if microcarriers consisting of other materials, such as gelatin, cellulose, plastic, or glass, are used. It is important to set up an appropriate fluorescence filter that considers the difference in fluorescence wavelength range between the microcarrier material and cells. Besides, some variability of our method may occur when the metabolic activity and redox state change because the total fluorescence intensity of autofluorescence varies with the state of NADH and FAD involved in metabolism and redox. To develop the more robust cell counting method based on our method, the degree to which the accuracy of the proposed method varies with differences in metabolic activity and redox state needs to be investigated in detail.

As one of the limitations of our method, it should be noted that this research only demonstrated the validity and feasibility of the proposed method as a proof-of-concept study at relatively low cell densities (10^3^-10^5^ cells/ml) for the required cell density of the industrial-level cultured meat technology (10^6^ cells/ml). Since cell counting seems to become more difficult at higher cell densities due to the signal saturation problem as with other label-free methods, the proposed method was validated in this study at cell densities of 10^6^ cells/ml or lower to demonstrate the proof of concept of the autofluorescence-based label-free cell counting method. If the method is to be used as an industrial-level technology in the cultured meat industry in the future, the validity of our method would need to be investigated at cell densities of 10^6^ cells/ml or higher. As another limitation of the proposed method, the noise that occur during the spatiotemporal averaging has an impact on the performance of the proposed method. As shown in Supplementary Fig. S2, it was observed that the curve of each statistic shifted to the positive direction of the x-axis when the noise was added. This result indicates that the required fluorescence intensity ratio may increase depending on the intensity of the noise. Therefore, it is necessary to consider setting an appropriate filter that increases the fluorescence intensity ratio and an appropriate number of shots when our method is applicated in a noisy condition.

In summary, we propose an inexpensive and robust label-free cell counting technique, that consists of a general-purpose fluorescence microscope and calculation of spatiotemporal average values. As the autofluorescence phenomenon is universal to cells, this technique is expected to contribute not only to the field of cultured meat but also to regenerative medicine and drug discovery, which require the mass culture of cells.

## Methods

### Concepts of our proposed method

In our proposed method, spatiotemporal data of the fluorescent signal are obtained by continuously capturing signals using a two-dimensional imaging device. The spatiotemporal average value is calculated through temporal averaging of the spatial mean pixel values at each time point (Fig. 1). Detailed information of the theory behind the proposed method is described in the Supplementary Note 1.

### Numerical experiments

To evaluate the effects of the fluorescence intensity ratio and number of shots on the proposed method numerically, we performed the Monte Carlo method based on the equation (3) and (5) of the Supplementary Note 1. The geometry of cells and microcarriers was set as spheres with diameters of 20 μm and 200 μm, respectively. The number of microcarriers and cells was assumed to follow a uniform distribution, and their fluorescence intensities were assumed to follow a normal distribution. Numerical experiments were conducted using Python 3.7.13 on an Intel Core i5-8250U CPU. For the Monte Carlo method, the permuted congruential generator was used as a pseudo-random number generator. Performances of the proposed method were quantitatively evaluated using correlation coefficient, coefficient of determination, absolute mean error, and root mean square error values. The coefficient of determination was calculated through regression analysis using a simple linear regression model. To reduce computation time, the noise term in equation (4) of the Supplementary Note 1 was set to ε ≪ 1. Other parameters used are listed in Supplementary Table S1.

To evaluate the effect of the fluorescence intensity ratio on the proposed method, we evaluated performance change with respect to the fluorescence intensity ratio ρ. The number of shots was set to 60 to simulate a system that measures fluorescence intensity at intervals of 1 s for 1 min. The fluorescence intensity ratios were set on a logarithmic scale from 10^−3^−10^3^.

To evaluate the effect of the number of shots on the proposed method, we assessed performance change according to the number of shots. The fluorescence intensity ratio was set on a logarithmic scale from 10^−2^−10^2^, and the number of shots on an integer scale from 1−60. To shorten the computation time, the process of calculating spatiotemporal average was processed using Cython 0.29.30, and the iterative computation was performed through multi-process parallel processing.

### Cell culture experiments

To evaluate the validity of the proposed method experimentally, we evaluated the performance of the proposed method by cell culture experiments. This study focused on cultured meat technology as a response to the recent severe food crisis [30] and protein supply shortage [31]. For this purpose, bovine myoblasts were used in the cell culture experiments. Bovine myoblasts were isolated as previously described [32, 33] and frozen after one passage. Dulbecco’s modified Eagle’s medium (043-30085; Fujifilm Wako Pure Chemicals) supplemented with 10% fetal bovine serum (10270-106; Gibco) and 1% Penicillin-Streptomycin-Amphotericin B Suspension (161-23181; Fujifilm Wako Pure Chemicals) was used as the culture medium. After thawing from frozen vials, bovine myoblasts were subcultured once and further cultured for four days on ϕ100 culture dishes (353003, Corning Inc.) coated with laminin-511. The laminin-511 coating was accomplished by spreading Easy iMatrix-511 silk (892024; Matrixome) on the bottom of the dish at a concentration of 0.06 mg/cm^2^ and incubating at 37 °C for at least 1 h. To prepare the laminin-511 coated microcarriers, a mixture of 30 mg polystyrene spherical microcarriers (Synthemax II, 3781; Corning) and 1 mL Easy iMatrix-511 silk solution was dropped in 12 well plates (HydroCell; CellSeed) and incubated for 1 h. After preparation of the laminin-511 coated microcarriers, the cell suspension with culture medium was mixed with the microcarriers in each well. Concentration of the final cell suspension was adjusted to 1, 2, 4, 6, 8, 10, 12, 14, 16, 18, 20, and 22-fold dilutions based on a maximum initial cell density of 5.00 × 10^5^ cells/mL. The number of cells was measured using our proposed method and the golden standard method after two days of culture.

Autofluorescence micrograph were captured using a fluorescence microscope (ECLIPSE Ti2; Nikon) equipped with a CMOS camera (DS-Qi2; Nikon). The exposure time was set to 5 ms and analog gain to 1×. An LED lighting system (D-LEDI, Nikon) was used as the light source, always at 100% illumination intensity. For fluorescence filters, DAPI (excitation: 371–409 nm, emission: 430–520 nm), FITC (excitation: 450–490 nm, emission: 507–562 nm), and TRITC (excitation: 525–575 nm, emission: 593–668 nm) filters were used. Sixty-one images were captured with each filter, and randomly sampled sixty images were used to analyze by the proposed method. To evaluate the reliability of our method, the bootstrap sampling with sixty iterations was performed. The images were saved in 14-bit grayscale using image measurement software (NIS-Elements BR version 5.30.03; Nikon), and the spatiotemporal averaging value was calculated using Python. During calculation, the arithmetic mean of the pixel values for an image was considered the spatial averaging value, and the arithmetic mean of the spatial means of multiple images was considered the spatiotemporal averaging value.

In the golden standard method, cells were collected using trypsin treatment and centrifugation, and the number of cells was measured using a hemocytometer. To evaluate the performance of the proposed method, a Bayesian linear regression between our method and the golden standard method was performed using No-U-turn sampler which is known as one of the efficient algorithms of Markov chain Monte Carlo simulation. The simulation was performed by a lightweight probabilistic programming library NumPyro version 0.11.0 [34].The number of samples generated by the simulation was 10^5^ to converge sufficiently. When *x* is the spatiotemporal average value measured by the proposed method, *y* is the cell density measured by the golden standard method and ε is the noise during the measurements, it can be formulated as *y* = *ax* + *b* + ε. Using the probabilistic simulation, the parameters *a, b*, and ε can be estimated with uncertainty. Given the range of cell densities in the culture experiments, a normal distribution with mean zero and standard deviation 10^5^ was used for the prior distribution of these parameters. After the estimation, performance of the proposed method was quantitatively evaluated using correlation coefficient, coefficient of determination, mean absolute error, and root mean square error values.

### Statistical analysis

The significance of the correlations was statistically evaluated using an uncorrelated test with Pearson’s correlation coefficient. *P* < 0.01 was accepted as statistically significant.

## Data availability

Data supporting the findings of this study are available in the article and Supplementary information files, or from the corresponding author upon request. All data generated and analyzed during this study are included in this published article (and its Supplementary Information file).

## Acknowledgements

We are grateful to Prof. Hironobu TAKAHSHI and Ms. Azumi YOSHIDA for isolation and preparation of the primary bovine cells and helpful discussions. This work was supported by the Japan Science and Technology Agency [grant number JPMJMI20C1] and partially by the Japan Society for the Promotion of Science KAKENHI program [grant number JP22K20417].

## Author Contributions Statement

T.M. and K.S. designed this study. T.M. fabricated the experimental system, conducted numerical experiment and cell culture experiment, analyzed the results, and wrote original draft. R.T provided cell resources, visualized results. K.S, R.T., K.I., and T.S discussed the results and supervised this study. T.M, K.S and T.S acquired the funding for this study. T.M., K.S., R.T., K.I. and T.S reviewed and wrote the manuscript.

## References

1. Bodiou, V., Moutsatsou, P. & Post, M.J. Microcarriers for upscaling cultured meat production. Front. Nutr. 7, 10; 10.3389/fnut.2020.00010 (2020).

2. Dabiri, S.M.H. et al. Multifunctional thermoresponsive microcarriers for high-throughput cell culture and enzyme-free cell harvesting. Small 17, e2103192; 10.1002/smll.202103192 (2021).

3. Treich, N. Cultured meat: Promises and challenges. Environ. Resour. Econ. (Dordr.) 79, 33–61; 10.1007/s10640-021-00551-3 (2021).

4. Tanaka, R.I., Sakaguchi, K., Yoshida, A., Takahashi, H., Haraguchi, Y. & Shimizu, T. Production of scaffold-free cell-based meat using cell sheet technology. npj Sci. Food 6, 41; 10.1038/s41538-022-00155-1 (2022).

5. Morikura, T., Sakaguchi, K., Tanaka, R. I, Yoshida, A., Takahashi, H., Iwasaki, K. & Shimizu, T. Conditioned serum-free culture medium accomplishes adhesion and proliferation of bovine myogenic cells on uncoated dishes. npj Sci Food 8, 108; 10.1038/s41538-024-00355-x (2024)

6. Ornelas-González, A., González-González, M. & Rito-Palomares M. Microcarrier-based stem cell bioprocessing: GMP-grade culture challenges and future trends for regenerative medicine. Crit. Rev. Biotechnol. 41, 1081–1095; 10.1080/07388551.2021.1898328 (2021).

7. Yang, J., Guertin, P., Jia, G., Lv, Z., Yang, H. & Ju, D. Large-scale microcarrier culture of HEK293T cells and Vero cells in single-use bioreactors. AMB Express. 9, 1–14; 10.1186/s13568-019-0794-5 (2019).

8. Knittel, J. et al. A microcarrier-based protocol for scalable generation and purification of human induced pluripotent stem cell-derived neurons and astrocytes. STAR Protoc. 3, 101632: 10.1016/j.xpro.2022.101632 (2022).

9. Shah, D., Naciri, M., Clee, P., Rubeai, M. & Rubeai, M. NucleoCounter – An efficient technique for the determination of cell number and viability in animal cell culture processes. Cytotechnology 51, 39–44; 10.1007/s10616-006-9012-9 (2006).

10. He, M.Y.C., Stacker, S.A., Rossi, R. & Halford, M.M. Counting nuclei released from microcarrier-based cultures using pro-fluorescent nucleic acid stains and volumetric flow cytometry. BioTechniques. 63, 34–36; 10.2144/000114568 (2017).

11. Dixit, R. & Cyr, R. Cell damage and reactive oxygen species production induced by fluorescence microscopy: Effect on mitosis and guidelines for non-invasive fluorescence microscopy. Plant J. 36, 280–290; 10.1046/j.1365-313x.2003.01868.x (2003).

12. Coutu, D.L. & Schroeder, T. Probing cellular processes by long-term live imaging - Historic problems and current solutions. J. Cell Sci. 126, 3805–3815; 10.1242/jcs.118349 (2013).

13. Degouys, V. et al. Dielectric spectroscopy of mammalian cells. Cytotechnology 13, 195–202. 10.1007/bf00749815 (1993).

14. Carvell, J.P. & Dowd, J.E. On-line measurements and control of viable cell density in cell culture manufacturing processes using radio-frequency impedance. Cytotechnology 50, 35–48; 10.1007/s10616-005-3974-x (2006).

15. Gong, L., Petchakup, C., Shi, P., Tan, P.L., Tan, L.P. & Tay, C.Y. Direct and label-free cell status monitoring of spheroids and microcarriers using microfluidic impedance cytometry. Small 17, e2007500; 10.1002/smll.202007500 (2021).

16. Margis, R. & Borojevic, R. Quantification of attached cells in tissue culture plates and on microcarriers. Anal. Biochem. 181, 209–211; 10.1016/0003-2697(89)90230-3 (1989).

17. Aijaz, A., Trawinski, D., McKirgan, S. & Parekkadan, B. Non-invasive cell counting of adherent, suspended and encapsulated mammalian cells using optical density. BioTechniques. 68, 35–40; 10.2144/btn-2019-0052 (2020).

18. Farrell, C.J. et al. Cell confluency analysis on microcarriers by micro-flow imaging. Cytotechnology 68, 2469–2478; 10.1007/s10616-016-9967-0 (2016).

19. Odeleye, A.O.O., Castillo-Avila, S., Boon, M., Martin, H. & Coopman, K. Development of an optical system for the non-invasive tracking of stem cell growth on microcarriers. Biotechnol. Bioeng. 114, 2032–2042; 10.1002/bit.26328 (2017).

20. Aubin, J.E. Autofluorescence of viable cultured mammalian cells. J. Histochem. Cytochem. 27, 36–43; 10.1177/27.1.220325 (1979).

21. Andersson, H., Baechi, T., Hoechl, M. & Richter, C. Autofluorescence of living cells. J. Microsc. 191, 1–7; 10.1046/j.1365-2818.1998.00347.x (1998).

22. Monici, M. Cell and tissue autofluorescence research and diagnostic applications. Biotechnol. Annu. Rev. 11, 227–256; 10.1016/s1387-2656(05)11007-2 (2005).

23. Surre, J., Saint-Ruf, C., Collin, V., Orenga, S., Ramjeet, M. & Matic, I. Strong increase in the autofluorescence of cells signals struggle for survival. Sci. Rep. 8, 1–14; 10.1038/s41598-018-30623-2 (2018).

24. Neumann, M. & Gabel, D. Simple method for reduction of autofluorescence in fluorescence microscopy. J. Histochem. Cytochem. 50, 437–439; 10.1177/002215540205000315 (2002).

25. Viegas, M.S., Martins, T.C., Seco, F. & do Carmo, A. An improved and cost-effective methodology for the reduction of autofluorescence in direct immunofluorescence studies on formalin-fixed paraffin-embedded tissues. Eur. J. Histochem. 51, 59–66; 10.4081/1013 (2007).

26. Torkelson, J.M., Lipsky, S., Tirrell, M. & Tirrell, D.A. Fluorescence and absorbance of polystyrene in dilute and semidilute solutions. Macromolecules 16, 326–330; 10.1021/ma00236a031 (1983).

27. Chakraborty, S., Harris, K. & Huang, M. Photoluminescence properties of polystyrene-hosted fluorophore thin films. AIP Advances 6, 125113; 10.1063/1.4972989 (2016).

28. Young, E.W.K., Berthier, E. & Beebe, D.J. Assessment of enhanced autofluorescence and impact on cell microscopy for microfabricated thermoplastic devices. Anal. Chem. 85, 44–49; 10.1021/ac3034773 (2013).

29. Awasthi, K., Chang, F.L., Hsieh, P.Y., Hsu, H.Y. & Ohta, N. Characterization of endogenous fluorescence in nonsmall lung cancerous cells: A comparison with nonmalignant lung normal cells. J. Biophotonics. 13, e201960210: 10.1002/jbio.201960210 (2020).

30. van Dijk, M., Morley, T., Rau, M.L. & Saghai, Y. A meta-analysis of projected global food demand and population at risk of hunger for the period 2010-2050. Nat. Food. 2, 494–501; 10.1038/s43016-021-00322-9 (2021).

31. Cole, M.B., Augustin, M.A., Robertson, M.J. & Manners, J.M. The science of food security. npj Sci. Food 2, 14; 10.1038/s41538-018-0021-9 (2018).

32. Takahashi, H., Yoshida, A., Gao, B., Yamanaka, K. & Shimizu, T. Harvest of quality-controlled bovine myogenic cells and biomimetic bovine muscle tissue engineering for sustainable meat production. Biomaterials 287, 121649; 10.1016/j.biomaterials.2022.121649 (2022).

33. Phan, D., Pradhan, N. & Jankowiak, M. Composable effects for flexible and accelerated probabilistic programming in NumPyro. Preprint at https://arxiv.org/abs/1912.11554 (2019)

