## Supplementary Information for "Autofluorescence-based Label-free Cell Counting Method in Suspension Culture with Microcarriers"

Takashi MORIKURA, Katsuhisa SAKAGUCHI*,

Ryu-ichiro TANAKA, Kiyotaka IWASAKI, Tatsuya SHIMIZU

*Corresponding author

**Supplementary Note 1**

Total fluorescence signal observed in the suspension culture system is measured as the sum of the fluorescence of cells and microcarriers. When a random variable of fluorescence intensity per unit volume of each cell at the excitation wavelength $\lambda$ is defined as $i_C \sim N(\mu_C, {\sigma_C}^2)$ and that of each microcarrier $i_M \sim N(\mu_M, {\sigma_M}^2)$, the spatial average value in a region of interest can be expressed using the following equation:

$$i_s = \frac{1}{L^2}\left\{\sum_{p=0}^{m-1}\left(v_M \cdot i_M + \sum_{q=0}^{n-1}(v_C \cdot i_C)_q\right)_p + \varepsilon\right\} \tag{1}$$

where $v_M$ and $v_C$ indicate the volume of microcarriers and cells, respectively. In addition, $m$ and $n$ indicate the number of microcarriers and cells in the test region, respectively, $L$ the length of one side of the region of interest, and $\varepsilon \sim N(0, {\sigma_\varepsilon}^2)$ the noise. In the Equation (1), it is assumed that multiple cells cover a single microcarrier, and the signal from the superposition of both is integrated for each microcarrier. When the spatial averaging value at time $t$ is represented by $(i_s)_t$, the spatiotemporal average value $I$ is expressed as follows:

$$I = \frac{1}{K}\sum_{k=0}^{K-1}(i_s)_{t=t_k} \tag{2}$$

In the equation (2), $K$ indicates the number of shots and $t_0$ indicate the initial time of the measurement indicates, respectively. Substituting equation (1) with equation (2), the space-time average value $I$ is expressed as follows:

$$I = \frac{1}{K}\sum_{k=0}^{K-1}\left[\frac{1}{L^2}\left\{\sum_{p=1}^{m-1}\left(v_M \cdot i_M + \sum_{q=1}^{n-1}(v_C \cdot i_C)_q\right)_p + \varepsilon\right\}\right]_{t=t_k} \tag{3}$$

This equation (3) indicates that the spatiotemporal average value $I$ is calculated by temporal averaging after calculating the spatial averages of fluorescence intensity to each image. Expanding equation (3) gives the following equation:

$$I = \frac{1}{KL^2}\left\{\sum_{k=0}^{K-1}\sum_{p=0}^{m-1}(v_M \cdot i_M)_{p,t=t_k} + \sum_{k=0}^{K-1}\sum_{p=0}^{m-1}\sum_{q=0}^{n-1}(v_C \cdot i_C)_{q,p,t=t_k} + \sum_{k=0}^{K-1}\varepsilon_{t=t_k}\right\} \tag{4}$$

If we consider the spatiotemporal average value $I$ as a random variable $I_K$, the reproducibility of the normal distribution and equation (4) would allow us to obtain the following results:

$$I_K \sim N(\mu_I,\ {\sigma_I}^2)\ where\ \ \mu_I = F(m) + \rho \cdot G(m,n), \rho = \frac{\mu_C}{\mu_M} \tag{5}$$

In the equation (5), the functions $F$ and $G$ indicate linearly coupled functions and ${\sigma_I}^2$ the variance of the spatiotemporal average value. In addition, $\rho$ indicates the fluorescence intensity ratio between the fluorescence intensity per unit volume of cells and microcarriers. Using equation (5), the following formula would hold for any $\epsilon > 0$ based on the weak law of large numbers.

$$\lim_{K\to\infty}\left(P(|\bar{I}_K - \mu_I| < \epsilon)\right) = 1 \; where \; \bar{I}_K = \frac{1}{K}(I_0 + I_1 + \cdots + I_{K-1}) \qquad (6)$$

where $\bar{I}_K$ indicates the average value of the random variable $I_K$. Equation (6) indicates that when the number of shots is increased the spatiotemporal average value approaches $\mu_I$, that is, $F(m) + \rho \cdot G(m, n)$. Therefore, if we could find any excitation wavelength band where the fluorescence intensity ratio $\rho$ increases, the spatiotemporal average value could be correlated to the number of cells under a sufficient number of shots.

In summary, by securing the number of shots under the specific excitation wavelength band where the fluorescence intensity ratio increases, a spatiotemporal average value would asymptotically correlate to the number of cells; hence, we hypothesized that the number of cells on microcarriers could be measured using the spatiotemporal average value.

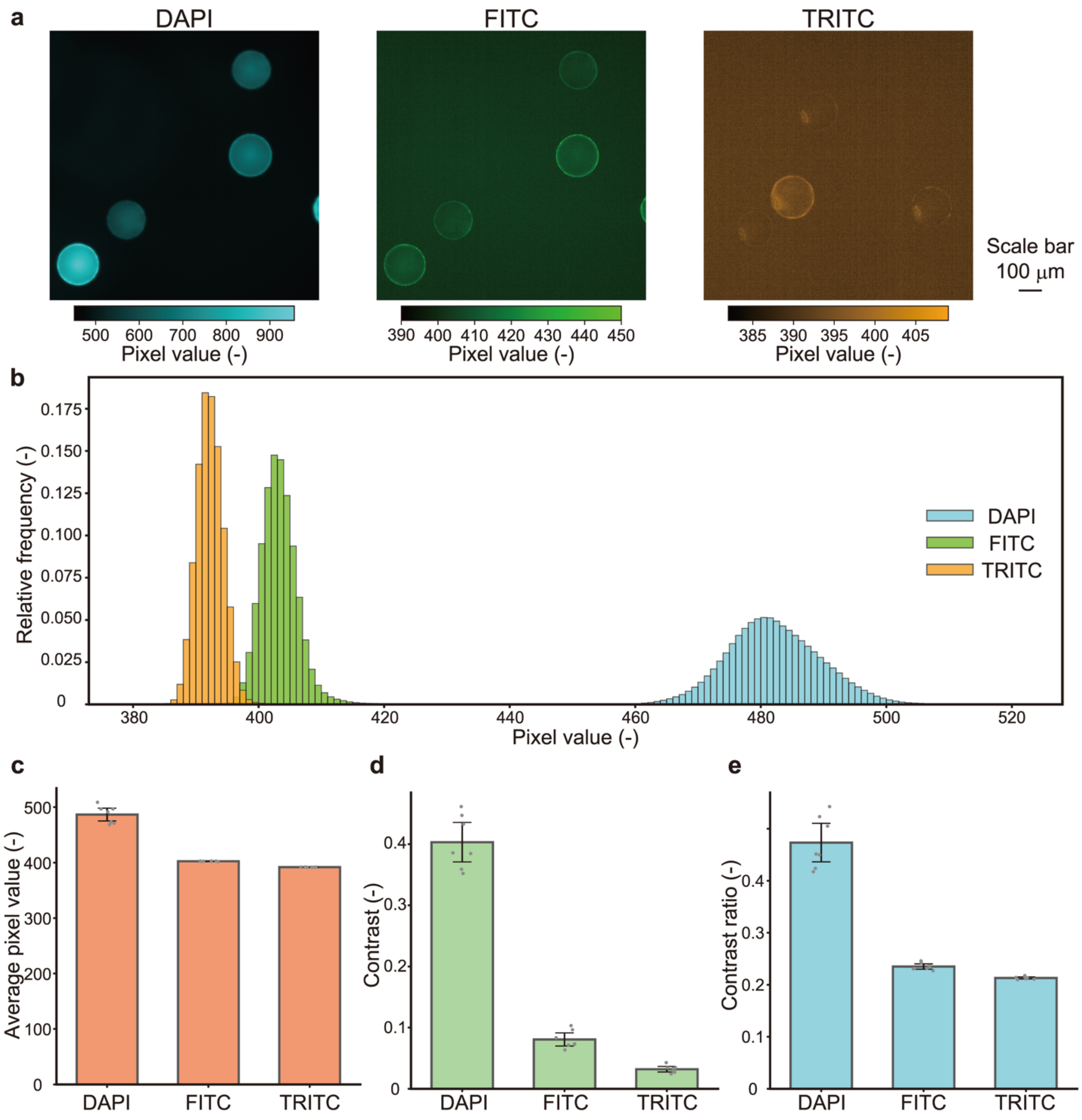


**Figure S1. Evaluation of the photoluminescence caused by Synthemax® II microcarriers.** The microcarriers (20 mg) were suspended in 5 mL Milli-Q water. Using DAPI, FITC, and TRITC filters, the microcarriers were irradiated with exposure times of 20 ms. (a) Representative fluorescence micrographs for each filter. (b) Representative histograms of the pixel value. Domain of the histogram was set to pixel values between 380 and 520. Relative frequency was calculated by normalizing the sum of the frequencies in each filter to 1. (c) Average pixel value of the image. (d) Image contrast. (e) Image contrast ratio. Sample size in this experiment was seven (mean ± 95% CI).

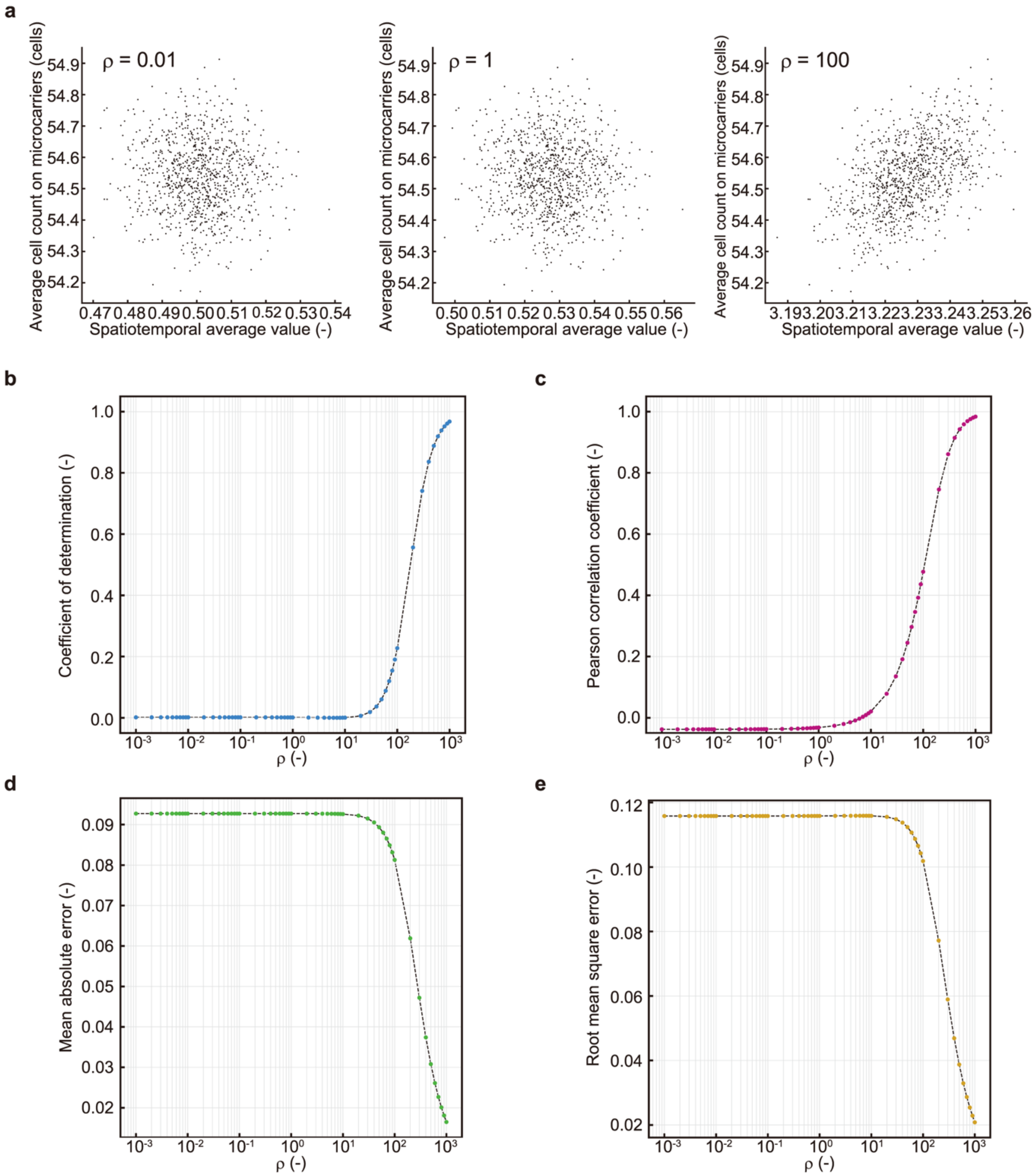


**Figure S2. Result of the numerical experiments with noise on the spatiotemporal average values.** The noise was set to be followed a normal distribution with mean 0 and standard deviation 0.01, which was 1/10 times of the minimum order of the spatiotemporal average value computed in the numerical experiment. Representative scatter plots of the average cell count on microcarriers and the spatiotemporal average value. The ratio of fluorescence intensity is indicated by ρ. Dots in the figures indicate samples generated using the Monte Carlo method. The number of samples was 1000. In addition, Results of numerical experiments on performance of the proposed method for the ratio of fluorescence intensity ρ were shown. (**a**) Coefficient of determination. (**b**) Pearson correlation coefficient. (**c**) Mean absolute error. (**d**) Root mean square error. Dots in the figures indicate samples generated using the Monte Carlo method. The number of samples was 1000.

**Table S1. Parameters used in numerical experiments.** Each value was determined based on the geometric constraints and preliminary experimental results. Specifically, the test area was assumed to be a square with approximately 1 cm on each side. The number of cells was determined based on the result of the preliminary experiments that the number of cells attached to the hemispherical surface was several dozen.

| | |
|---|---|
| Maximum number of microcarriers | 500 |
| Minimum number of microcarriers | 100 |
| Maximum number of cells | 80 |
| Minimum number of cells | 30 |
| Fluorescence intensity per unit volume of microcarriers | 0.0005 |
| Coefficient of variation of the fluorescence intensity distribution of microcarriers | 10 |
| Coefficient of variation of the fluorescence intensity distribution of cells | 10 |
